# On Negative Heritability and Negative Estimates of Heritability

**DOI:** 10.1101/232843

**Authors:** David Steinsaltz, Andy Dahl, Kenneth W. Wachter

## Abstract

We consider the problem of interpreting negative maximum likelihood estimates of heritability that sometimes arise from popular statistical models of additive genetic variation. These may result from random noise acting on estimates of genuinely positive heritability, but we argue that they may also arise from misspecification of the standard additive mechanism that is supposed to justify the statistical procedure. Researchers should be open to the possibility that negative heritability estimates could reflect a real physical feature of the biological process from which the data were sampled.

## 1. Introduction: The meaning of heritability

### 1.1. Operational definitions of heritability

As Albert Jacquard [15] pointed out decades ago, *narrow-sense heritability* — commonly denoted *h*^2^ — has conventionally two distinct meanings:

1. The proportion of total variance attributable to additive genetic effects;
2. The slope of the linear regression of children’s phenotypes on the mean parental phenotypes.

Both meanings appear in the earliest works to give a quantitative operational definition to *heritability*, in particular [21]. (For more about the history of the notion of heritability see [2].)

The correspondence between these two meanings depends on an additive model, where genetic and non-genetic effects are independent and sum together to produce the phenotype. When we have general genetic relatedness (rather than parental relations with fixed 50% relatedness) heritability is analogous to a regression coefficient relating phenotypic similarity to genotypic similarity.

We are particularly concerned here with the interpretation of negative estimates of heritability. The appearance of negative estimates for a parameter of crucial scientific interest that is *prima facie* positive is unusual, as has often been noted. Negative estimates of the heritability parameter are often dismissed as a mathematical abstraction, values in a range that arises purely formally and that may only be reported for formal purposes, as part of an ensemble of estimates that collectively are unbiased. Several recent studies [32, 30, 3, 7, 34, 11, 10, 14] have reported individual negative heritability estimates in this way, including them in averages that themselves came out positive. We illustrate one such analysis [17] using RNA-sequencing data from the GEUVADIS consortium in section 5.

In [16] a point estimate of −0.109 is obtained for heritability of horn length in Soay sheep. It is immediately dismissed with the statement that “it is impossible to have negative heritability” and the inference is drawn that the true heritability must actually be a small positive number toward the upper end of the confidence interval.

We wish to argue that negative heritability estimates need to be taken more seriously. The confusion, we contend, comes from the overlap between statistical models that operationalize the two different interpretations of heritability described above. The argument for rejecting negative estimates appears compelling just so long as the focus is only on the random-effects probability model in equation (1) that typically motivates our definition 1 of heritability. Variance is nonnegative, hence the ratio of two variances cannot be negative. The denominator represents total variance and the numerator represents one component of variance, implying a ratio in [0,1] if the two components are independent (as the model presumes).

While “variance attributable to additive genetic effects” is a basic element of the genetic model, it has no place in the statistical algorithms such as GREML derived from this model that is widely used to estimate heritability from experimental data. The GREML algorithm is actually (as we will explain in section 2.1) the realization of a multivariate normal model that is naturally constrained to have the parameter 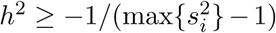, where 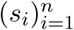 are the singular values of the genotype matrix. If the phenotypes were derived from summing independent additive genetic effects then the true *h*^2^ must indeed be nonnegative, but that must be recognized as an assumption that must be scientifically warranted, as it is not compelled on formal grounds.

In contemporary analyses of genomic data, two roles played by alleles may be distinguished. Alleles exercise or tag actual causal influences on traits. Alleles also serve as markers of family and ancestry, markers of relatedness among individuals that may structure historical, behavioral, social, and environmental influences on traits. As attention expands beyond the additive genetic model to data-analytic genome-wide statistical studies, heritability and the parameter called heritability are not necessarily the same.

### 1.2. The meaning of negative heritability

Once we have accepted the GREML multivariate normal framework — which we will define precisely — we must admit the possibility that the joint distribution of phenotypes and genotypes in a given dataset may be best described by an *h*^2^ value that is negative. The question this raises is, can such a negative heritability estimate be biologically sensible? As described in Section 2, the parameter for heritability may be identified, in a precise way, with a correlation between genotype similarity and phenotype similarity. The model invites us to select an estimate of *h*^2^ that will best match the genetic covariance between individuals to the similarity in their traits. Even if we *want* heritability to be interpreted in the first sense, as a partition of variance, this will not, in general, be correct. All we have access to from the data is an estimate of something like heritability in sense 2. High heritability means that individuals with similar genotype tend to have similar trait values. Zero heritability means that genotypes tell us nothing about similarities in trait values. Negative heritability, then, could be perfectly sensible as a description of the data: It means that individuals with similar genotypes are likely to have more divergent trait values than those with highly disparate genotypes.

Saying that a given set of data might be best described by a negative heritability estimate goes only part of the way toward answering the question of the biological plausibility of the concept. Suppose you were estimating the weight of water droplets by successively adding them to a small container, and estimating the slope from the sequence of weights. If the scale is sufficiently imprecise it is hardly unlikely that we could estimate a negative slope, yet common sense tells us that negative estimates should be dismissed as unrealistic, and truncated at zero (or combined with prior expectations to form a positive Bayesian estimate). Statistical theory tells us that this system should produce slope estimates that may be positive or negative, but that the probability of a negative estimate goes to zero as the number of measurements goes to infinity, and the estimate converges to its true (positive) value. The essential question is, is there a plausible mechanism that could produce genuine negative heritability, so that as the amount of data generated by the model goes to infinity, the estimate converges to a negative quantity.

The term “negative heritability” appeared for the first time, so far as we are aware, in a paper [12] by J. B. S. Haldane, written around 1960, but first published posthumously in 1996. Haldane described how the maternal-effect trait of neonatal jaundice could be said to display negative heritability: Because the disease results from maternal antibodies against a fetal antigen, it will not arise in a fetus whose mother herself experienced neonatal jaundice^1^. Haldane then calculates a negative heritability from a model that is specialized to the peculiar inheritance structure of neonatal jaundice.

We will suggest one such mechanism in Section 4. As with Haldane’s model (which may be understood as a special case), this mechanism has implications which may be implausible or even obviously false in a given experimental setting. It involves interactions between individuals that are not primarily genetic, and so may be dismissed as irrelevant to the study of genetic heritability. The point we want to suggest, though, is that as an abstract physical mechanism that could be producing our data it is as mathematically plausible as the linear random-effects model that undergirds GREML. This is only one example of such a mechanism, and the conclusion we advocate is that negative heritability must be acknowledged as a genuine phenomenon for genotype-phenotype data, even if it may be reasonably excluded by the context of some particular studies. Thinking about what sorts of biological settings could yield negative heritability can also prove an effective guide to understanding when negative heritability estimates may be reliably truncated or ignored.

This is very much like the advice on “interpretation of negative components of variance” propounded in a very different context by the statistician J. A. Nelder [22] in 1954. Nelder considered the problem of ANOVA testing on split-plot experiments, where error for main plots was found to be smaller than the error for subplots, producing a negative estimate for the residual subplot error. As we have done here, Nelder showed how the apparently negative “variance component” could arise either from sampling error from a positive variance component, or from a misspecification of the model, where correlations between measurements have been neglected. “In any particular situation,” Nelder concludes, “it is the statistician’s responsibility to decide which model is more appropriate.”

## 2. The GREML model as linear regression

### 2.1. The random-effects model

For the remainder of this paper we follow [26] in using the letter *ψ* to represent heritability, to avoid the confusing implication built in to the nomenclature *h*^2^ that this parameter cannot be negative.

Underlying GREML, as well as alternative approaches to heritability estimation such as LD-score and Haseman-Elston regression, is a basic random-effects model. Following the notation of [26], our basic object is a data set consisting of an *n × p* matrix *Z*, taken to represent the genotypes of *n* individuals, measured at *p* different loci. There is a vector **y**, representing a scalar observation for each of the *n* individuals. The underlying observations are counts of alleles taking the values 0, 1, or 2, but the genotype matrix is centered to have mean zero in each column and normalized to have mean square over the whole matrix equal to 1. (Often, columns are further standardized to variance 1, but we do not make this assumption). The model posits the existence of a random vector **u** ∈ ℝ*^p^* of genetic influences from the individual SNPs such that

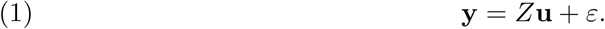

The vectors **u** and *ε* are assumed to be independent and to have zero means and i.i.d. normal components. The variances are determined by two parameters, which are to be estimated: *θ* represents the precision (reciprocal variance) of the non-genetic noise and *ψ* represents the heritability, entering the model as the ratio of genetic variance to total variance. We will also use the notation *φ* = *ψ*/(1 — *ψ*) in some places, for concision.

The GREML model has been formulated as a random-effects model, but it is equivalent to a multivariate normal model corresponding to the covariance matrix

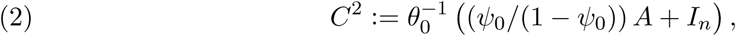
where *A* = *ZZ*^*^/*p* is the Genetic Relatedness Matrix (GRM), and *θ*_0_ and *ψ*_0_ are the true values of the parameters. In this section we describe how the model may also be understood as a linear regression model.

In their original paper [31], Yang and coauthors spell out an analogy between GCTA and a different form of linear regression. They regress squared trait differences between pairs of individuals on corresponding elements of the GRM, with *n*(*n* − 1)/2 points and correlated errors (this is Haseman-Elston regression, which has recently become a popular heritability estimation method due to its speed and robustness to some degree of model misspecification [9, 5]). Instead, we draw an approximate comparison between GREML and regression with *n* points and independent errors.

Let 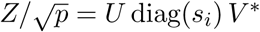 be the singular-value decomposition of 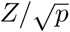, and rotate the observations to diagonalize the covariance matrix, obtaining

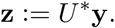

Because the columns of *Z* have zero means, one of the singular values is zero and the corresponding column of *U* is proportional to a vector with all elements equal to 1. Thus in each other (orthogonal) column of *U* the elements sum to zero, and each column defines a contrast between weighted groupings of individuals in the sample.

The elements of **z** are independent centered normal random variables, and *z_i_* has variance 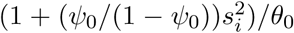. It follows that 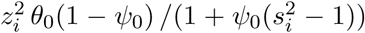 are independent chi-square random variables each on one degree of freedom and

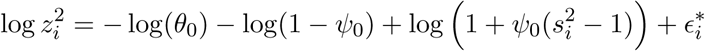
where the 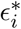 are distributed as the logarithms of the independent chi-square variables, long-tailed to the left, with mean ≈ −1.302, standard deviation ≈ 2.266, and skewness ≈ −1.643.

The mean of 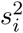 is 1, and when 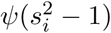 are uniformly small we may approximate our equation by

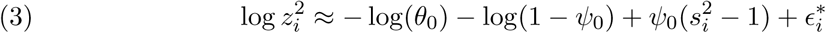

Here *ψ*_0_ takes on the role of the true slope for a regression of log 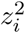 on 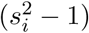. It can be estimated by least squares, bearing in mind that the skew of the 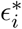 affects standard errors of estimation.

Practitioners of GREML instead usually estimate 0 via maximum likelihood. It is shown in [26] that the maximum likelihood estimates can be expressed in terms of quantities 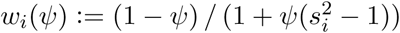 and 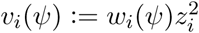. They satisfy

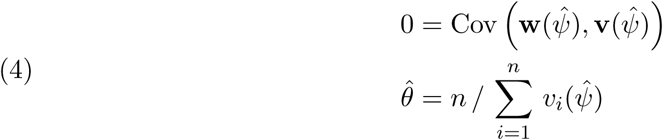

Here Cov is the empirical covariance of vector elements, an operation on vectors defined by 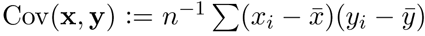, and Var is similarly defined. We also set *τ*_2_(*ψ*) = *ψ*^−2^Var(*w*(*ψ*)), and omit the dependence on *ψ* when helpful. When *ψ*_0_ is modest and the variance of the squared singular values is small, the least-squares and maximum likelihood estimates are close to each other.

Suppose, however, that the true variances of the *z_i_* include a phenotypic contribution that varies inversely with the singular values of *Z*. In the model developed in Section 4 to first order in 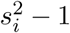 the true slope is 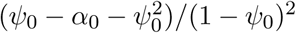 as a function of a repulsion parameter *α*. When *α*_0_ exceeds *ψ*_0_, the true slope turns negative and estimated slopes correctly include negative values. The precision of the estimated slopes depends on the variance of the squared singular values. When the 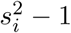 are tightly clustered, there will be large standard errors for the estimates of *ψ*.

### 2.2. Simulations

We illustrate in Figure 1 a close match between regression-based and maximum likelihood estimates for *ψ*. The plot also emphasizes the wide estimation uncertainty when the singular values 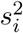 are tightly clustered.

**Figure 1.**
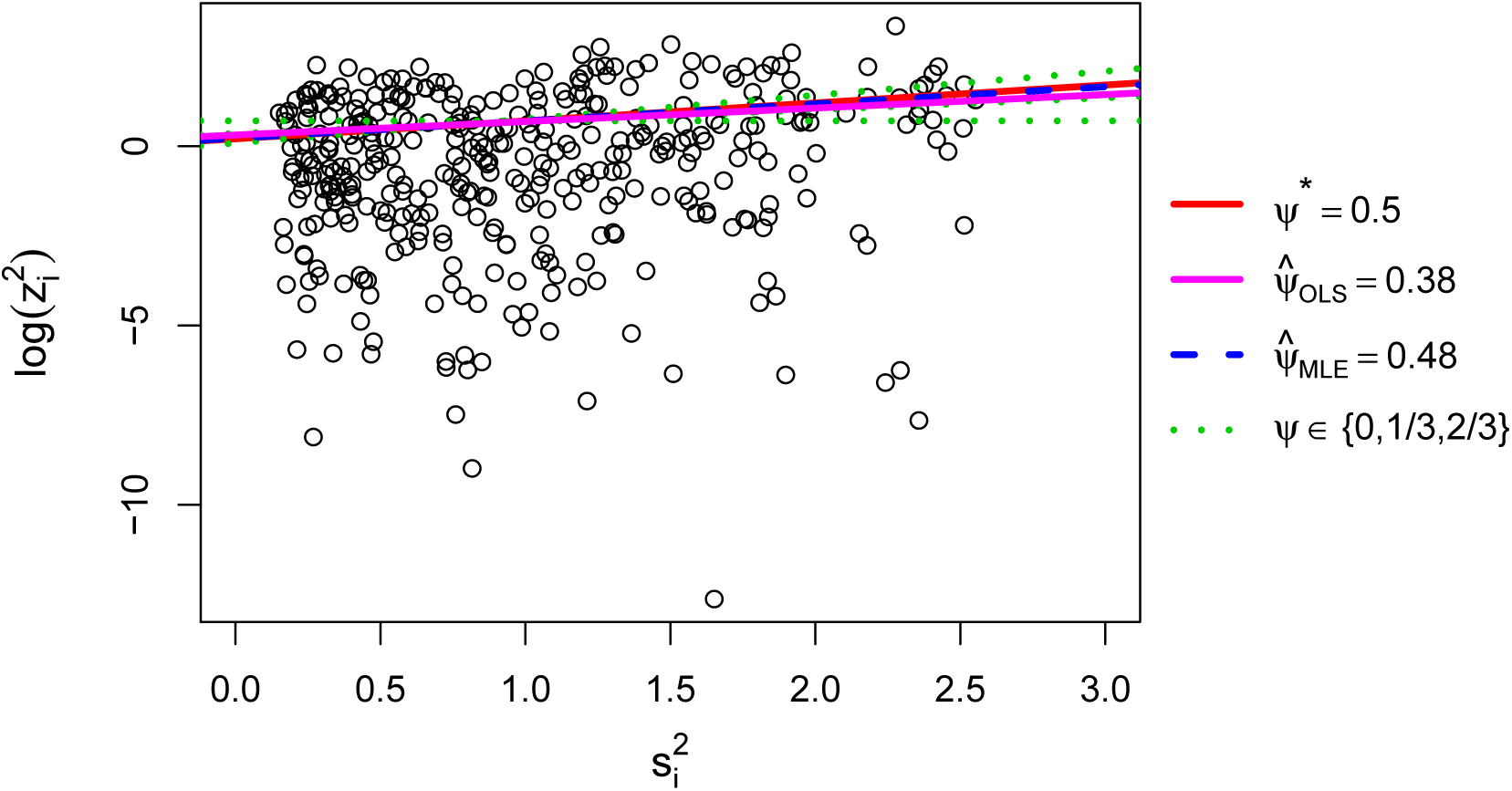
Linear-regression approximation of the estimation problem for *ψ*. The solid red line shows the true line, corresponding to *ψ* = 0.5, while the blue and pink lines respond to the MLE and linear regression estimator, respectively. The dotted green lines correspond to *ψ* = 0, 0.33, 0.67, 0.84.

We draw 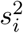 from the so-called independent setting, which assumes the genotype matrix *Z* has independent entries. The known limiting measure for these singular values is the Marcenko-Pastur distribution, from which we draw *n* equally-spaced values. We directly simulate the independent whitened phenotypes *z*_i_ as independent normal variables with mean zero and variance 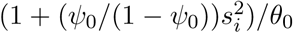. This procedure gives *ψ* estimates that are approximately equivalent to those obtainable from simulating a random *Z* with i.i.d. entries along with the associated random trait vector *y*. See section 5 of [26] for details.

We choose *Z* to have *n* = 375 rows and *p* = 1,000 columns, similar to the dimensions of the *cis*-window genotype matrices in Section 5. Nonetheless, the 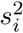 from our independent setting differ from those empirical singular value distributions. We could simulate a different model for the GRM simply by choosing a different distribution for 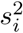. We also choose *ψ*^*^ = 0.50 and *θ*^*^ = 1.

To compute the MLE, we solve the first equation in (4), a simple univariate function of *ψ*, and substitute into the second equation. To compute our regression-based estimator, we perform ordinary least squares regression of 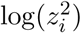 on 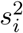 and an intercept.

Figure 1 shows a scatterplot of one realization of pairs 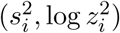. The red line corresponds to the true value of *ψ*, i.e. *ψ*^*^ = .5, and describes the expectation of 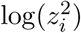 as a function of 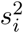 given *θ*^*^ = 1. The solid pink line corresponds to the ordinary least squares estimate and is very similar to the dashed blue line for the MLE. Dotted green lines are shown for several more distant alternative values for *ψ*. It is no wonder that slopes are hard to pin down in the face of such huge scatter in observations.

## 3. Bias from rejecting negative heritability estimates

The common practice of truncating the maximum likelihood calculation to non-negative values introduces bias that is well-known and may be serious for samples of moderate size, both when estimates are truncated at zero and when negatives are ignored. It is thus worth looking beyond the original motivation to the actual structure of the GREML model and considering what meaning negative parameter values might turn out to have.

The problem of estimating the probability of negative heritability estimates was studied fifty years ago by [8]. We add here a few comments about how the framework described in [26] may contribute to understanding the magnitude of the negative heritability estimate problem that arises from sampling noise in settings where the true heritability is understood to be nonnegative, hence where truncation at zero (or rejection of negative estimates) is warranted and guarantees improved estimates in, say, mean squared error. We gain a rough idea of the impact of rejecting negative estimates from a normal approximation

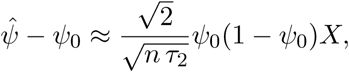
where *X* has standard normal distribution (see [26] for derivation). Rejecting estimates where 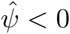, we have the conditioning bias

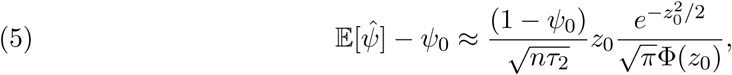
here 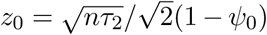 and Φ is the standard normal c.d.f. If we instead truncate the estimates — raising all negative 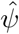 to 0 — we get the truncation bias

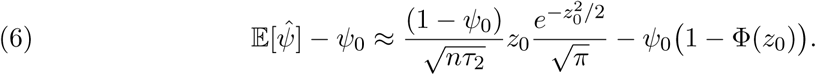

## 4. The phenotypic repulsion model

The notion that new species force their way into phenotypic gaps in the existing ecological community was termed by Darwin the “principle of divergence” and has been further developed by ecologists under the name “phenotypic repulsion” or “phylogenetic repulsion” [28]. Species living in close proximity — which are often closely related phylogenetically — coexist by separating from each other phenotypically. A similar kind of competitive exclusion has been proposed [27] on the individual level to explain observed pattern of developmental variation within human families. Social niche-formation within families has also been proposed by [13] — without an explicit mathematical model — as the basis for an evaluation of gene-environment interaction based on misclassified twin types. While we are not aware of mathematical models of this phenomenon, one could certainly imagine local competition for sunlight, combined with range-limited seed dispersion, yielding an effective phenotypic repulsion between related plants in a forest setting, or monozygotic twins who seek to distinguish themselves from their sibling.

We propose a model of phenotypic repulsion where individuals that are most closely related genetically strive to avoid each other phenotypically. We begin with a model like that described in Section 2.1, where individuals have phenotypes determined by normally distributed effect sizes acting on their individual genotypes. We introduce a penalty term to the probability, of the form

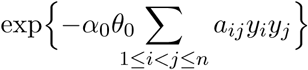
where 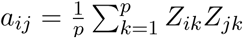 is the (*i*, *j*) entry of the GRM, and *α* ≤ 1 is a parameter with true value *α*_0_ measuring the extent to which genetically similar individuals are pushed to have differing phenotypes. Of course, this setup could be generalized to higher-dimensional phenotypes, with *y_i_y_j_* replaced by an arbitrary inner product. The penalty term is inspired by the statistical mechanics models that have been applied to geographically-structured population dynamics, such as the Contact Process [20], used to model the spread of epidemics.

Combining this specification with (2) we see that the phenotypes will now be multivariate normal with mean 0 and covariance matrix

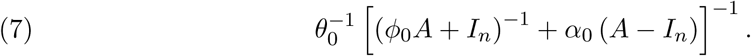

It follows that the transformed phenotypes **z** = *U*^*^**y** are independent normal with mean zero and variance

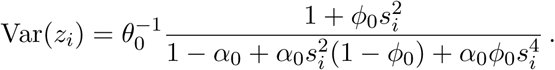

Suppose the data have come from this phenotypic-repulsion model, and we analyze them using the random-effects model. While it is always possible to get 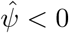 because of random fluctuations, we would like to show that the heritability implied by this model is “really” negative, in the sense that the distribution of *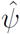* converges to a strictly negative value as the number of subjects goes to ∞. This will follow from Proposition 4.1 (below) with

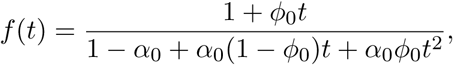
as long as *φ*_0_ < *α*_0_, since

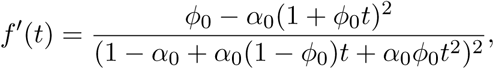
which is less than 0 for all *t ≥* 0.

In other words, to the extent that we say that heritability is defined by the linear model, heritability can be negative if genotypes and phenotypes interact through the environment in a manner like the phenotypic repulsion model. This will be true even if the phenotypic interactions are limited to small family groups. We prove that this is the case — that the heritability to which the estimates converge with increasing population size is negative — in the following Proposition, which is proved in section ??.

### Proposition 4.1.

*Suppose we have a family of n* × *n GRMs A_n_ for n → ∞*, *with eigen-values 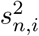. We suppose that the distributions of eigenvalues converge to a nontrivial distribution dσ*(*s*^2^).

*Let U*^(^*^n^*^)^ *be the corresponding eigenvector matrix. For each n we have a multivariate normal random vector* **y**^(^*^n^*^)^ *with covariance matrix U*(*^n^*) 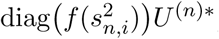, *where f*: ℝ^+^ → ℝ^+^ *is a strictly decreasing, continuously differentiable function. We assume that the singular values s*_max_ *are bounded above by s*_max_, *and that*

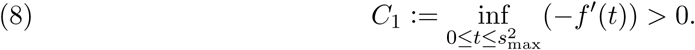

*(We maintain the normalization assumption that 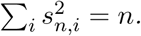)*

*Let 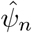 be the MLE for an observation* **y**^(^*^n^*^)^, *calculated from the random-effects model with GRM A_n_. Then 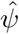 is bounded above in probability by a strictly negative quantity* −*δ*, *depending on C*_1_ *and the distribution σ, as n →* ∞. *That is, the probability of 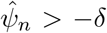 goes to 0 as n* → ∞.

## 5. Transcriptome-wide Heritability Estimation in GEUVADIS

We end on a brief example to illustrate the practical significance of negative heritability estimates. Although negative estimates of heritability for a single, fixed trait are rarely published, it is common to include negative estimates when profiling heritability across a large number of roughly exchangeable traits [32, 30, 3, 7, 34, 11, 10, 14]. Characterizing such -omic-wide heritability is common in functional genomics, where high-throughput measurements of some genomic property are made at thousands of genomic positions. The most common measurement is (RNA) gene expression, but other prominent examples include methylation levels, chromatin accessibility, expression response to stimuli, or protein expression.

We analyzed an RNA-sequencing dataset from the consortium on Genetic European Variation in Health and Disease (GEUVADIS) [17] ^2^. We aligned the raw transcript reads from the European individuals to the reference hg19 transcriptome with the RSEM software package that implements an Expectation-Maximization (EM) algorithm [19]. We removed perfectly correlated genes and genes with low expression mean or variance.

For each *i* in 375 people and *j* in 4,154 genes, we define the phenotype 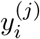 as log (1 + *n_ij_*) where *n_ij_* is the number of observed RNA reads for gene *j* measured in person *i*. Our preprocessing resulted in a sample for 4,154 genes measured on 375 people, each of which we centered and scaled to mean zero, variance one.

Separately for each gene *y*^(^*^j^*^)^, we estimate its *cis*-heritability, that is, the heritability in expression levels explained by SNPs near to the gene. We do so by fitting our standard model (1) with a genotype matrix *Z*^(^*^j^*^)^ whose columns correspond to SNPs located up to 1 megabase upstream or downstream of gene *j*’s transcription start site. Restricting to SNPs near a gene is a common way to enrich for functionally causal SNPs. We discard rare SNPs, which we define as SNPs with minor allele frequencies below 2.5%. Finally, we remove genes with less than 1,000 corresponding SNPs, which excludes 35 genes.

The column dimensions (p) of the separate genotype matrices range from 1, 000 to 20, 523 with a mean of 3,027 and median of 2, 754. We fit each *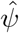* with the maximum likelihood routine from [6], yielding 4,119 values reflecting systematic variation across genes in their *cis*-heritability, within the limits imposed by sampling error.

The distribution of the resulting transcriptome-wide (*cis*-)heritability estimates is shown in Figure 2 in the form of a smoothed histogram. Clearly, a substantial number of the estimates are negative. The mode is close to zero. Removing negative heritability estimates increases the transcriptome-wide average heritability from 6.2% to 9.0%, and truncating at zero increases it from 6.2% to 6.6%.

**Figure 2.**
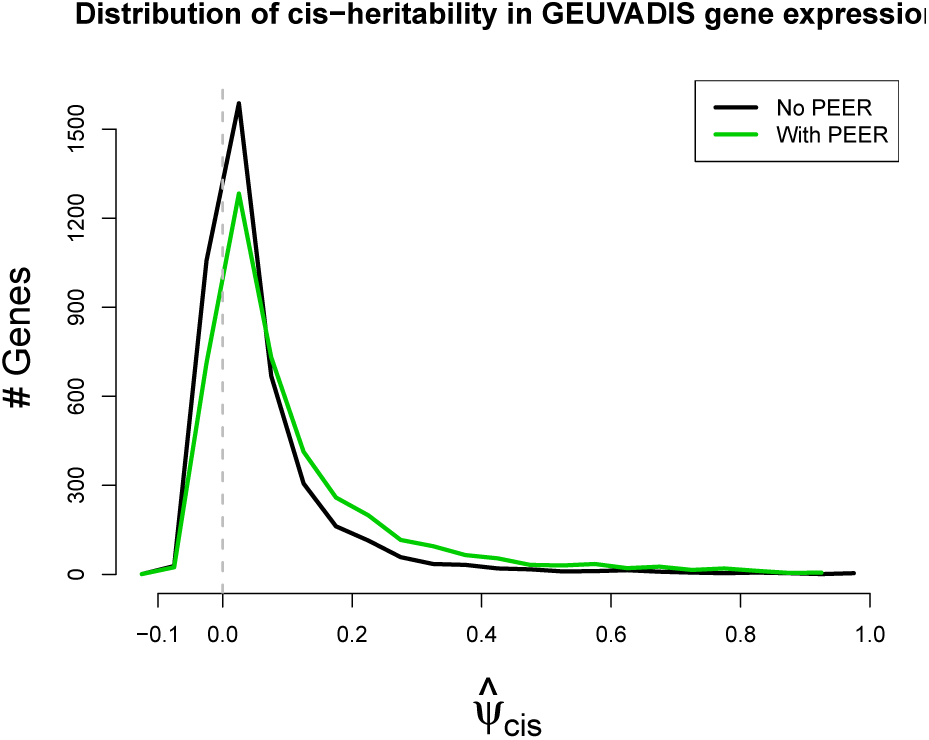
Transcriptome-wide density of gene expression *cis*-heritability estimates in the GEUVADIS data.

We repeated the analysis with adjustments based on the technique of Probabilistic Estimation of Expression Residuals (PEER) [24]. We projected out the top 10 PEER factors from the expression phenotypes and the SNPs in *Z*. This practice, or close variants based on gene expression principal components [1] or surrogate variables [18], is nearly universal in functional genomics [25]. The common aim of these approaches is to approximate latent confounding variation, like experimental batch effects, which can be captured by dimensionality reduction under certain assumptions. The confounder estimates are treated as known covariates and residualized from the phenotype and genotype data.

Correcting for 10 PEER factors increases many of the *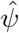* values and reduces the incidence of negative 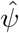 as shown in the green curve in Figure 2. However, it is clear that many negative estimates remain. Negative estimates are bound to be part of the picture whenever *ψ*_0_ is small and estimated with low precision, both conditions which will likely hold in most functional genomic analyses for at least the near future. On the question of whether some negative estimates may be meaningful reflections of non-genetic phenotypic structure, it is well to keep an open mind.

## 6. Proof of Proposition 4.1

### Proof.

We follow the general principle, enunciated by [29], that the MLE for the misspecified model will converge to the closest fit in the Kullback-Leibler sense. In other words, the parameter estimate converges in probability to the location of the maximum *expected value* of the log-likelihood function. The arguments of [29] do not apply directly here, because we are not sampling i.i.d. random variables; however, the score function may be written

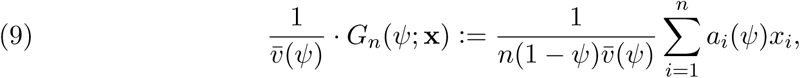
for 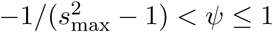, where (*x_i_*) are i.i.d. 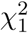 random variables and

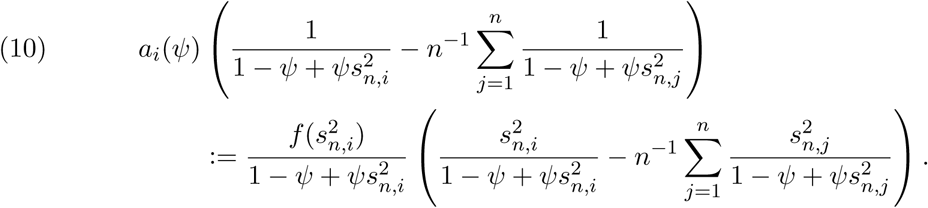

We note that the maximum likelihood occurs either at a zero of *G_n_*, or at *ψ* = 1 if *G_n_* is everywhere positive. (It goes to + ∞ at the left boundary.) We restrict to *ψ* ∈ [*β*_1_, *β*_2_] for some fixed *β*_1_ < *β*_2_ strictly in the interval 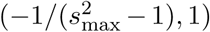. The coefficients *a_i_*(*ψ*) are uniformly bounded and uniformly Lipschitz, so, by a variant of the central result of [33], *G_n_*(*ψ*; **x**) converges uniformly in *ψ* to the function that is the limit of the expected values

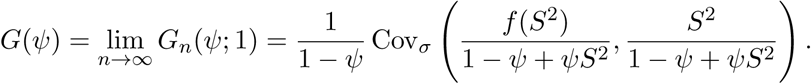

The covariance is understood here to be with respect to *S*^2^ having distribution *σ*. (This result does not satisfy exactly the conditions of [33], so we provide a proof of the result, stated as Lemma 6.1.) We have assumed that the limiting distribution *σ* for the squared singular values is nontrivial, hence with nonzero variance. We may assume without loss of generality that

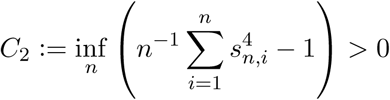

We need to show that *G* is negative for all *ψ* above the bound given in (11).

Near *ψ* = 0 the function *G_n_*(*ψ*; 1) is well-behaved, and takes on the value 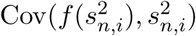 at *ψ* = 0. Since *f* (*t*) + *C*_1_*t* is a decreasing function of *t*, for 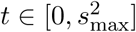, we have by Harris’s inequality [4, Theorem 2.15]

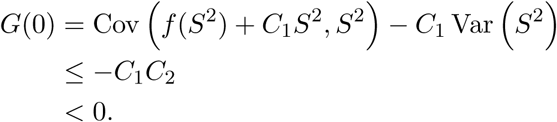

We also have

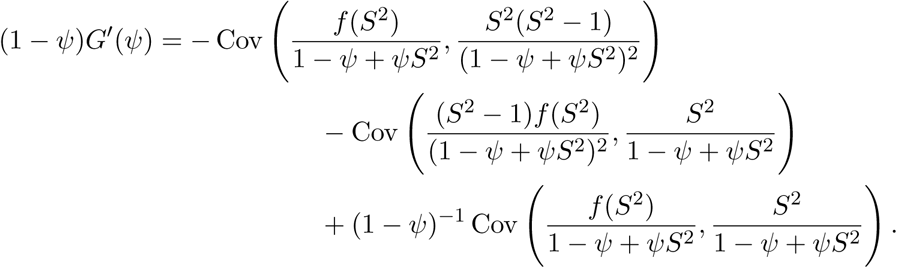

Since *f* is decreasing, we have for 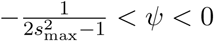 the bound

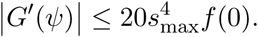

Thus *G*(*ψ*) < 0 for all

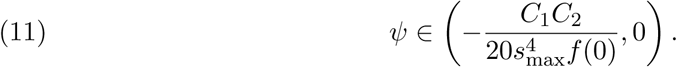

It follows that *G_n_*(*ψ*; **x**) is negative for all *ψ* between 0 and the bound given in (11), with probability tending to 1 as *n →* ∞.

We note now that for *ψ* ∈ [0, 1] *f*(*t*)/(1 − *ψ* + *ψt*) is a decreasing function of *t*, and *t*/(1 − *ψ* + *ψt*) is increasing, so (again by Harris’s Inequality) *G*(*ψ*) < 0, which completes the proof.□

### Lemma 6.1.

*Let* a: [0, *s*_max_] × [*β*_1_, *β*_2_] → ℝ *be a function that is bounded and Lipschitz. Let s_n_*_,1_.,…, *s_n,n_ be a triangular array of uniformly bounded real numbers, with a limiting measure σ, so that* 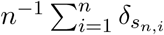 *converges weakly to σ, where δ_x_ represents unit point mass at x. Let x_n_*_,1_,…, *x_n,n_ be independent random variables with* 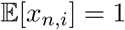 *and* Var(*x_n,i_*) ≤ *V for a fixed number V. Define*

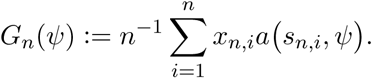

*Then G_n_ converges weakly in the uniform topology to*

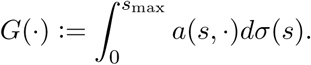

*That is*,

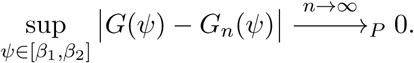

#### Proof.

We have

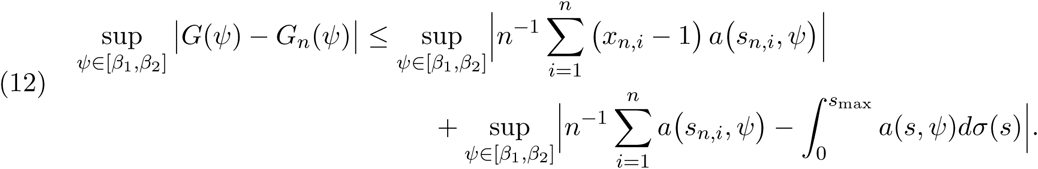

The Lipschitz property of *a* implies that the second term may be bounded, for any fixed positive integer *k*, by

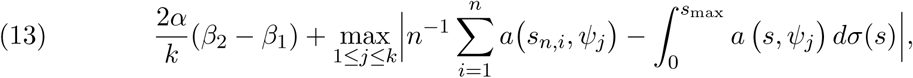
where

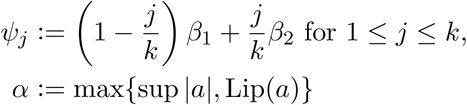

Because of the assumed weak convergence of the distribution of the *s_n,i_*, this converges to 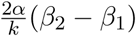 as *n* → ∞ for each fixed *k*; hence the second term converges to 0 as *n* → ∞.

To deal with the first term we use the standard method of *chaining (cf*. [23, chapter 3]): We define finite skeletons of [*β*_1_, *β*_2_], subsets *D*_0_ ⊂ D_1_ ⊂ · · · with |*D_j_*| = 2*^j^*, defined by

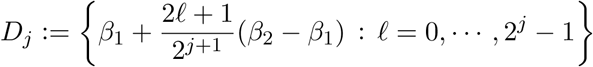

Note that each *ψ* ∈ *D_j_* for *j* ≥ 1 has a unique nearest neighbour in *D_j_*_−1_, which we will denote by *ψ*′, and |*ψ* − *ψ*′| = (*β*_2_ − *β*_1_)2^−^*^j^*^−1^. Define

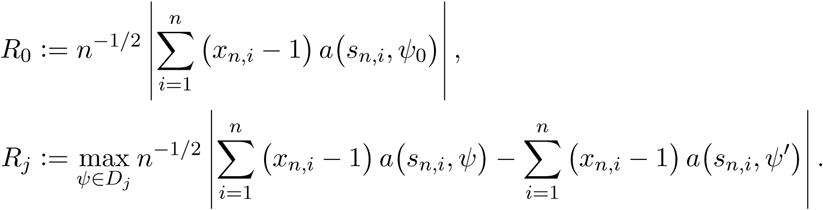

We have

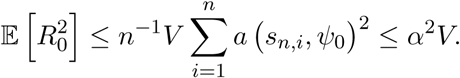

For any collection of random variables ξ_1_,…,ξ*_m_* we know that

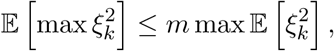
so

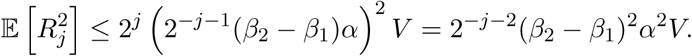

By the Cauchy–Schwarz inquality we have

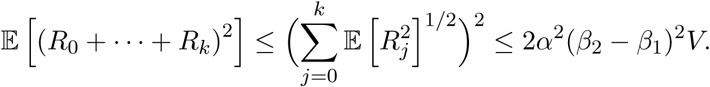

So finally, since

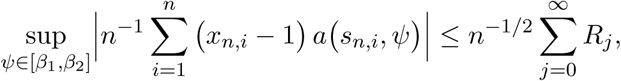
we have

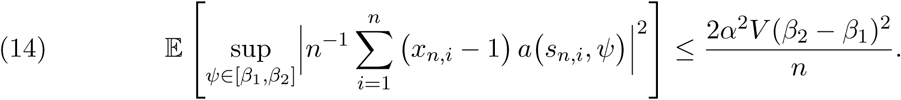

By Markov’s inequality, combining this with (13) completes the proof.□

## Footnotes

1 We thank Jonathan Marchini for pointing out this reference to us.

2 We thank David Siegel for help processing the GEUVADIS data.

## Notes

DS supported by Grant ES/N011856/1 from the UK Economic and Social Research Council. AD supported by Grant 1U01HG009080–01 from the U.S. National Institute of Health. KWW supported by Grant 5P30AG012839 from the U.S. National Institute on Aging.

